# Accurate modeling of replication rates in genome-wide association studies by accounting for Winner’s Curse and study-specific heterogeneity

**DOI:** 10.1101/856898

**Authors:** Jennifer Zou, Jinjing Zhou, Sarah Faller, Robert P Brown, Sriram S Sankararaman, Eleazar Eskin

## Abstract

Genome-wide association studies (GWAS) have identified thousands of genetic variants associated with complex human traits, but only a fraction of variants identified in discovery studies achieve significance in replication studies. Replication in GWAS has been well-studied in the context of Winner’s Curse, which is the inflation of effect size estimates for significant variants due to statistical chance. However, Winner’s Curse is often not sufficient to explain lack of replication. Another reason why studies fail to replicate is that there are fundamental differences between the discovery and replication studies. A confounding factor can create the appearance of a significant finding while actually being an artifact that will not replicate in future studies. We propose a statistical framework that utilizes GWAS and replication studies to jointly model Winner’s Curse and study-specific heterogeneity due to confounding factors. We apply this framework to 100 GWAS from the Human GWAS Catalog and observe that there is a large range in the level of estimated confounding. We demonstrate how this framework can be used to distinguish when studies fail to replicate due to statistical noise and when they fail due to confounding.

## Introduction

Replication is a gold standard in scientific discovery. Consensus emerges when a result has been replicated repeatedly by multiple researchers. Recently, a vigorous discussion has emerged of how often replication of an initial study fails across all fields of science, including genomics [1, 2, 3, 4, 5].

Genome-wide association studies (GWAS) are an ideal model to study replication because there are a large number of GWAS data sets with replication studies publicly available. GWAS replication studies are typically conducted in an independent cohort using only the variants that were significant in the initial (“discovery”) study. In the National Human Genome Research Institute Catalog of Published GWAS, thousands of genetic variants have been associated with complex human traits but not all associated variants achieve significance in replication studies [6, 4, 5, 7].

There are several reasons why associations do not replicate. The first is simply statistical. It is possible that the association is not observed in the replication study by chance. However, if the p-value from the original finding is highly significant and the replication studies have similar experimental designs, this scenario is unlikely. A second reason why studies can fail to replicate is Winner’s Curse, which is the inflation of effect size estimates for significant variants in a study due to statistical chance. This phenomenon occurs because the reported findings are a small fraction of many possible findings. In the case of GWAS, the significant associations are discovered after examining millions of variants and pass a stringent genome-wide significance threshold. This can result in inflated effect size estimates of significant variants in a study, especially when studies have low power [8]. Winner’s Curse has been studied extensively in GWAS, and multiple methods have been proposed to correct for its effects [9, 10, 11, 12, 4]. However, Winner’s Curse is often not sufficient to explain lack of replication. A third reason why studies fail to replicate is that there are fundamental differences between the discovery and the replication study or “study-specific heterogeneity”. An effect present in one study but not present in other studies can create the appearance of a significant finding that is not replicated in future studies [13]. This can either occur because of an underlying biological difference or a technical difference between the two studies. We refer to the cause of these differences as confounders.

Current methods for modeling confounders fall into two broad categories. The first class of methods attempts to model the effect of confounders before the association statistic is calculated in order to remove their effects from the association statistic [14, 15]. While these methods are widely used, they have several fundamental limitations. Methods that account for known covariates may not correct for all potential confounders. Confounding correction methods that use unsupervised learning to learn principal components or other global patterns in the data can incorrectly model the true signal as a confounder, which would remove true biological signal from the data [16, 17]. Similarly, when using unsupervised methods, it is unclear when there is residual confounding that remains in the data. The second class of methods attempts to directly adjust p-values by a constant factor to remove inflation [18, 19]. An example of such a method is genomic control [18]. In genomic control, there is an assumption that relatively few variants affect the trait. The implication of this assumption is that if the association statistics are ranked, then the variant corresponding to the median statistic will not affect the trait, and the value of this statistic will represent only the effect of the confounders. Genomic control scales all of the p-values using this statistic. Recently it has been observed that due to polygenicity and linkage disequilibrium (LD) structure in the genome, the majority of variants (including the one corresponding to the median statistic) either affect the trait or are correlated with variants that affect the trait. This breaks the genomic control assumption.

While LD-score regression has been shown to distinguish polygenicity and confounding [19], it has been shown that this approach can also result in inflated SNP-based heritability estimates under strong stratification [20].

In this paper, we present a novel approach for characterizing study-specific heterogeneity due to confounders using replication studies. The key insight in our approach is that we can use replications to estimate the effects of confounders and then account for their effects. Since replication studies are performed on the same phenotype, utilizing replication studies to estimate the effect of confounders does not rely on assumptions about the genetic architecture of the trait to distinguish between polygenicity and confounding. Furthermore, we can apply our approach in combination with traditional techniques like linear mixed models and regressing out the effect of covariates that are applied before computing association statistics. Our approach can be used to model any residual confounding effects after application of these methods.

In our framework, we perform a bivariate analysis between the z-scores from the discovery study and the z-scores from the replication study, while modeling the effects of both Winner’s Curse and study-specific confounders. We show through simulations that we can accurately estimate the contribution of study-specific confounders on a study and use this estimate to explain observed patterns of replication. We apply this framework to 100 GWAS from the Human GWAS Catalog and observe that there is a large range in the level of confounding observed across GWAS. We show that our estimate levels of confounding correlates well with observed patterns of replication and demonstrate how this can be used to distinguish when studies fail to replicate due to statistical noise and when they fail due to confounding.

## Results

### Method overview

The main goal of this framework is to account for Winner’s Curse and study-specific confounding in discovery and replication GWAS of the same phenotype. We compare this model to a naive model that only accounts for Winner’s Curse. We introduce these two models without accounting for difference in sample size for clarity, but we relax this constraint in the Methods section.

In GWAS, Winner’s Curse is the phenomenon where the association statistics for variants meeting a genome-wide threshold tend to be overestimated. Winner’s Curse can be observed in Figure 1, where the association statistics for the significant variants in the discovery study are substantially lower in the replication study. Due to this phenomenon, not all of the significant variants in the discovery study replicate. Winner’s Curse is widely observed in GWAS due to lack of statistical power in discovery studies. When power is low, the variants that are most significant in a study are likely to have inflated effect sizes due to random noise.

**Figure 1:**
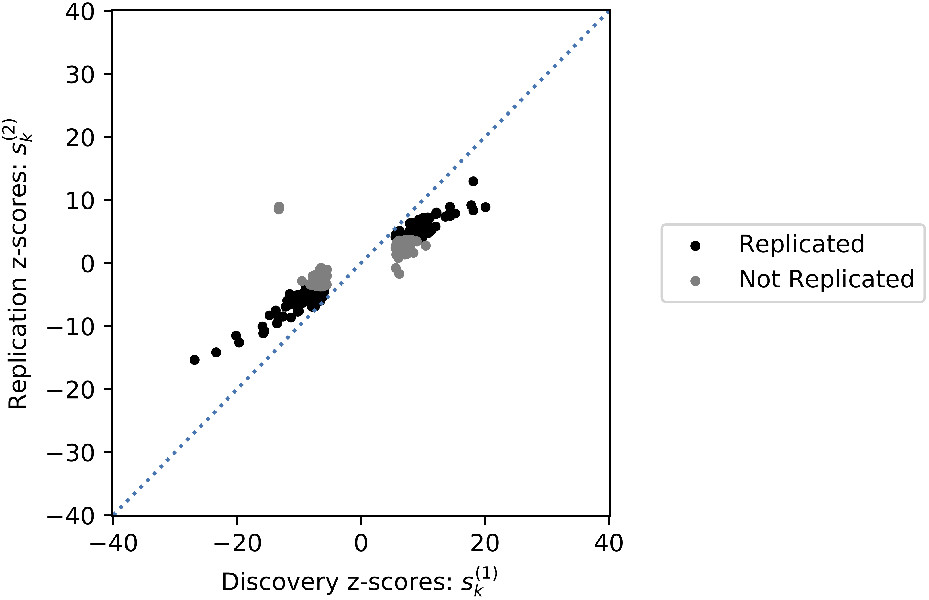
Bivariate GWAS analysis. We perform a bivariate analysis between the z-scores of the discovery and replicate GWAS (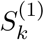and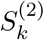, respectively). The significant variants in a discovery GWAS study on height (PMID 25282103) are shown. The variants that replicated successfully using a Bonferonni threshold of 0.05 are shown in black, and the ones that did not replicate are shown in grey. Many variants have stronger associations to the phenotype in the discovery study than the replication study. This phenomenon can be partially explained by Winner’s Curse and partially explained by study-specific confounders. This method jointly models the effects of Winner’s Curse and study-specific confounders on the observed z-scores.

To model random noise contributing to Winner’s Curse, we model the statistics for each variant *k* from the discovery and replication studies as normally distributed random variables (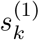 and 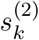, respectively). We assume that there is a shared genetic effect *λ* that is responsible for the observed association signal. Thus, the distribution of the statistic for variant *k* in study *i* is 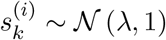. We define the prior probability of the true genetic effect to be 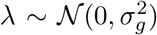, where the variance in the true genetic effect is learned from the data. Then, we model the joint distribution of the statistics from the two studies (Equation 1).

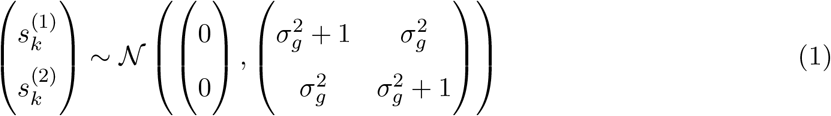

We correct for Winner’s Curse by computing the conditional distribution of the replication statistics given the discovery statistics (Equation 2). Using this conditional distribution, we can compute the expected value of the statistic in the replication study, along with confidence intervals on this estimate. This framework accurately models the data in cases where Winner’s Curse is the only source of inflation. Figure 2A shows a GWAS on height [21], where most of the variants fall within the 95% confidence intervals of the model accounting for Winner’s Curse. This shows that in studies without substantial confounding effects, Winner’s Curse can adequately explain the the proportion of variants that replicate or replication rate.

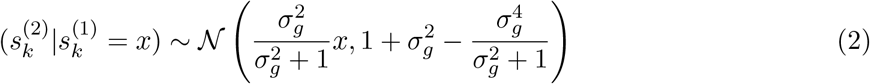

**Figure 2:**
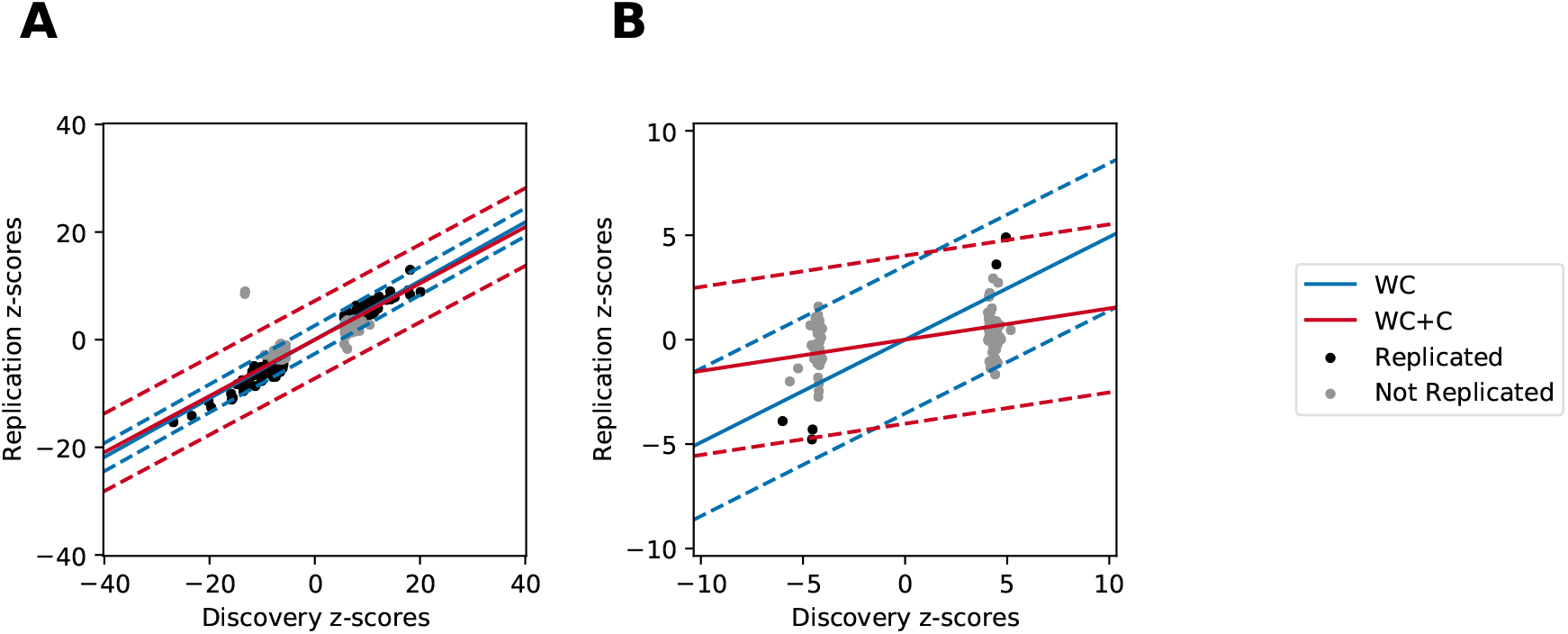
Correcting for Winner’s Curse and confounding. The x-axis for each plot is the value of the discovery z-score, and the y-axis is the value of the replication z-score. The solid lines correspond to the expected values of the replication z-score given the discovery z-score. The dotted lines represent confidence intervals in the estimates. The blue lines correspond to the model that only accounts for Winner’s Curse (WC), and the red lines correspond to the model that accounts for Winner’s Curse and confounding (WC+C). A) In this GWAS on height (PMID 25282103), there is very little confounding, and a model that accounts for Winner’s Curse explains the majority of the data. B. In this GWAS on height in African American women (PMID 22021425), there is substantial confounding. The model accounting for only Winner’s Curse (blue) does not explain the observed data well, whereas the model with Winner’s Curse and confounding (red) does explain the data well.

However, there is often additional heterogeneity due to confounding, where a framework that only accounts for Winner’s Curse would not explain the data well. Figure 2B shows an example of a GWAS on height in African American women [22]. In this study there is substantial confounding, and only 18% of variants replicate. Using a model that only accounts for Winner’s Curse, most variants are outside of the 95% confidence intervals, indicating that there is additional heterogeneity that is not modeled. To account for study-specific confounding, we decompose the effect size of the statistics into a genetic effect (*λ*) and study-specific confounding effects (*δ* ^(*i*)^). The distribution of the statistic for variant *k* in study *i* is 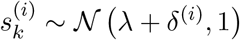. In addition to the prior on the genetic effect, we introduce priors on the study-specific confounders 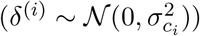.We incorporate both of these priors into the joint distribution of the statistics (Equation 3).

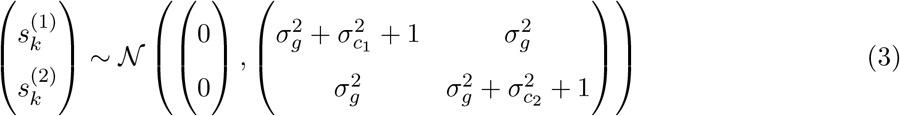

We correct for both Winner’s Curse and confounding by computing the conditional distribution of the replication statistic given the discovery statistic (Equation 4). By taking into account the additional variance in the association statistics from confounders, we are able to more accurately model the statistics from the two studies (Figure 2B). We quantitatively assess how well each model fits the data by estimating the number of variants that should replicate under each model (Methods). The naive model that only accounts for Winner’s Curse estimated that 56% of variants would replicate, whereas our model that also accounts for confounding estimated that 18% of variants would replicate, which is closer to the observed replication. This difference in the estimated replication under each model is due to the study-specific confounding effects estimated in the second model, which both decreases the expected value of the statistics in the replication study and increases the variance of the statistics in the replication study. After correcting for Winner’s Curse and confounding, most variants are within the 95% confidence intervals for the model. Thus, in this study, modeling study-specific confounders is necessary to explain the observed patterns of replication.

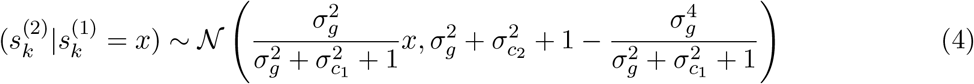

We apply this framework to simulated data and 100 human GWAS in the GWAS catalog. For all data sets, we compute the expected replication rate under the two models in order to compare how well each model fits the data, relative to each other.

### Winner’s Curse and confounding accurately explains replication in simulated data

To demonstrate that our approach accurately models the effects of Winner’s Curse and confounding to explain replication rates, we generated simulated data, where the variance in the genetic 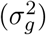 and confounding effects 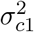 and 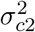 are known. In all simulations, we set the sample size of the discovery study to be 2000 and the sample size of the replication study to be 1000. We simulated z-scores for 1 million independent variants in each simulation.

For each simulation, we fixed 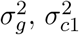, and 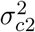 to be one of four values and simulated true effect sizes and study-specific confounding effects for each variant. We then simulated z-scores for the two studies (Methods). For each combination of parameter values, we repeated this 1000 times to generate a total of 64,000 simulations. We then used a Bonferonni corrected significance threshold of 5e-8 to identify variants that were significant in the discovery study. The number of significant variants in the discovery study for each simulation ranges from 263-11,362 variants (Figure S1A). We computed the replication rates as the proportion of variants in the discovery study that met a nominal threshold of 0.05 in the discovery study and had the same direction of effect in the two studies. The observed replication ranged from 15% - 60%. On average, higher levels of 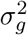, yielded higher replication rates (Figure S1B).

For each simulation, we used the z-scores of variants significant in the discovery study and their corresponding z-scores in the replication study as input to our model. We computed the maximum likelihood estimates (MLE) of 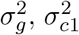, and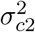. The estimates of the parameters were accurate and unbiased for all simulations (Figure 3).

**Figure 3:**
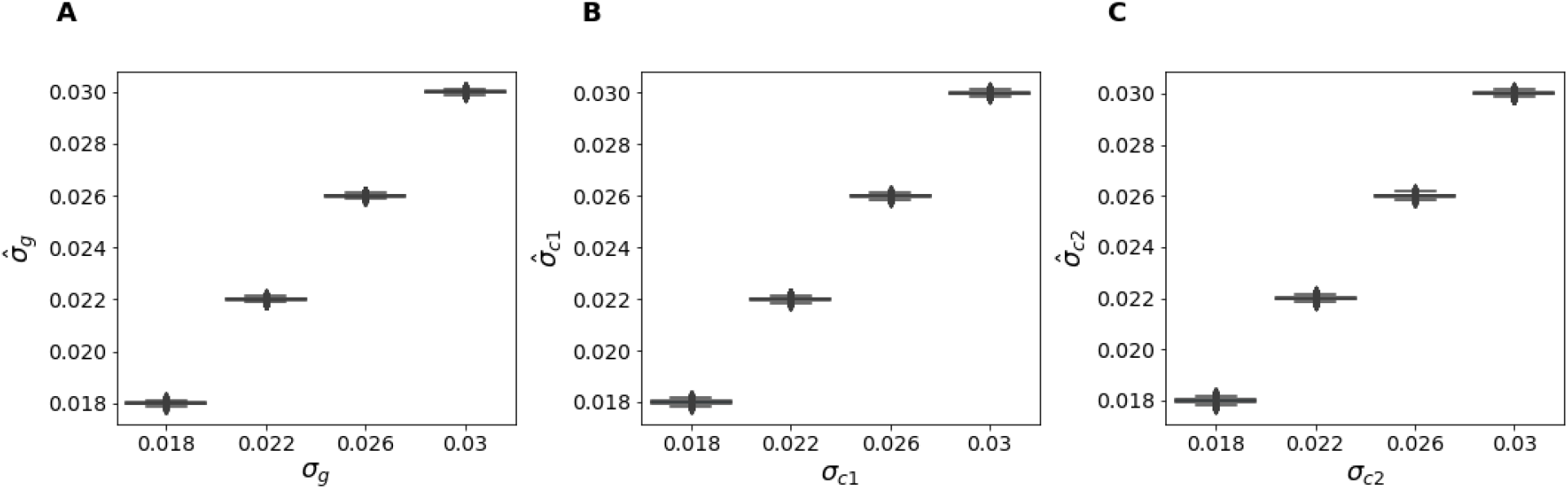
Variance components in Winner’s Curse and Confounding Simulations. True values of variance components (x-axis) vs estimated values (y-axis) for A) 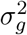 B) 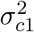 C) 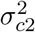

We used the MLE parameters to compute the expected replication rate under the model and compared this to the observed replication rate to assess the fit of each model. The WC+C more accurately described replication compared to the WC model (Figure 4A and Figure 4B), which we would expect since the simulations have study-specific confounding. Additionally, the expected replication rate using the MLE parameters are close to the expected replication rates using the true values of the parameters, indicating that the expected replication rate is robust to variance in the parameter estimates (Figure S2).

**Figure 4:**
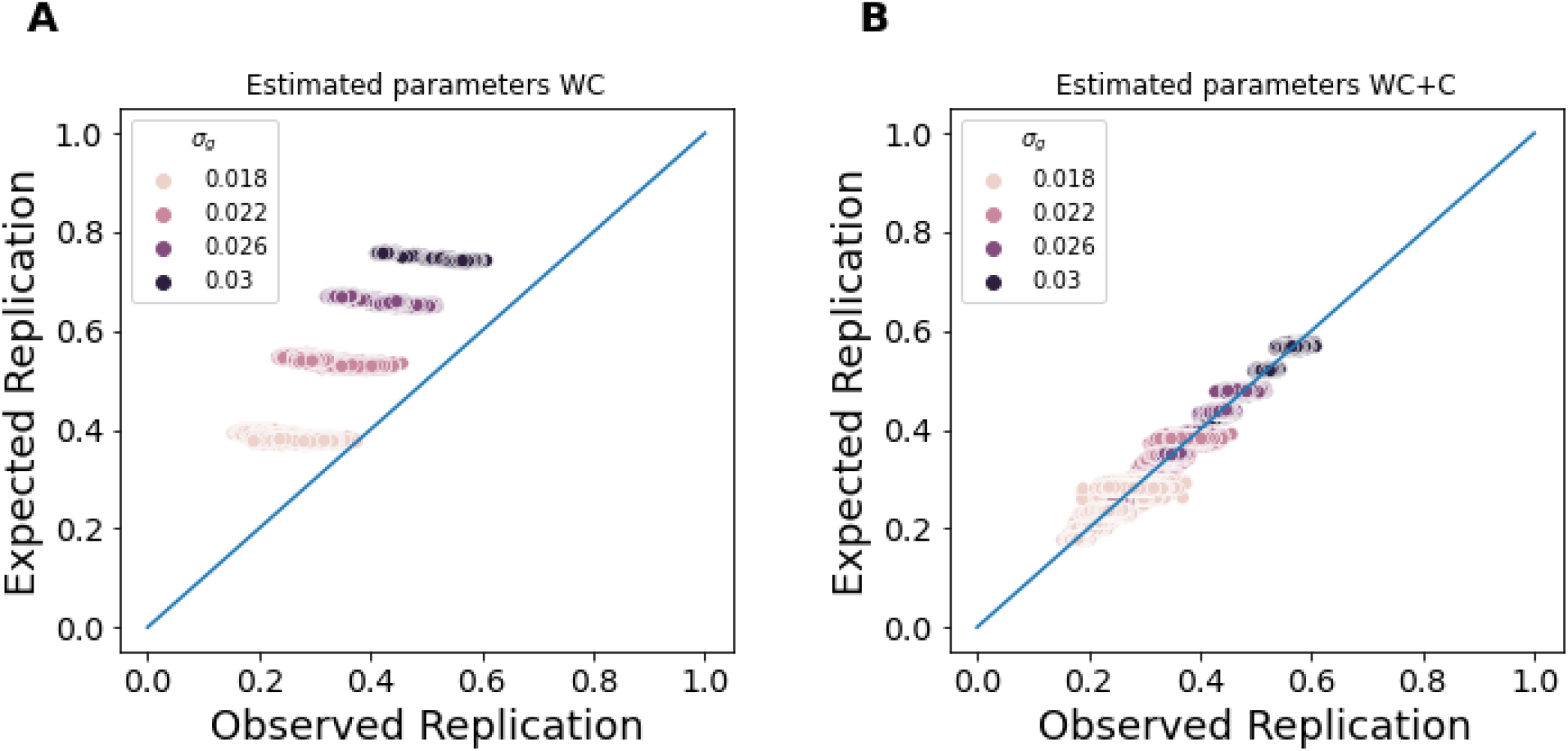
Winner’s Curse and Confounding Simulations. A) We computed the expected replication rate under the WC model. The WC model over-estimates replication because it does not account for confounding between the studies. B) We computed the expected replication rate under the WC+C model. The WC+C model more accurately describes the observed replication than the WC model.

### Accounting for missing data in discovery and replication designs

In studies with discovery and replication designs, often only a subset of variants are tested in the replication study. In some studies, only the summary statistics for variants that were significant in the discovery study are reported. The variants that were not significant in the discovery study are missing data. For these missing variants, we compute the likelihood of the data by integrating over all possible values of the data given the significance threshold used in the discovery study (Equation 19).

To evaluate whether the MLE estimates of the parameters are accurate in these situations with missing data, we used the previous set of simulations, where the variance in the genetic 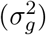 and confounding effects (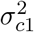 and 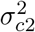) are known. For this set of analyses, we used the z-scores of the significant variants in the discovery study and their corresponding z-scores in the replication study only to estimate the parameters. The estimates of the parameters were accurate, but the variance in the parameter estimates was higher (Figure S4). Despite the higher variance in the parameter estimates, the expected replication rate using the MLE parameters are close to the expected replication rates using the true values of the parameters, indicating that the expected replication rate is robust to variance in the parameter estimates (Figure S3). Similar to previous simulations, the WC+C more accurately described replication compared to the WC model (Figure 5A and Figure 5B).

**Figure 5:**
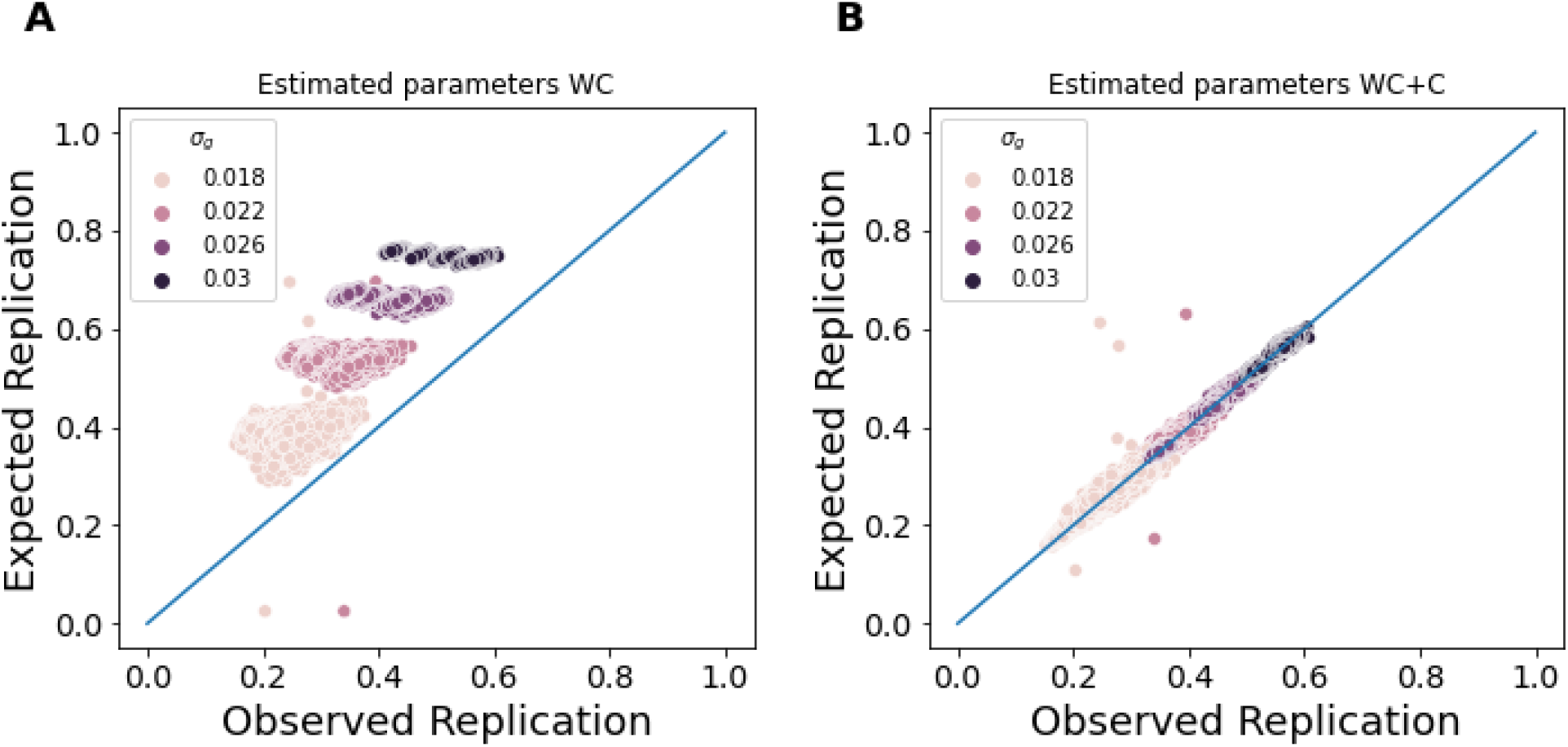
Winner’s Curse and Confounding Simulations. A) We computed the expected replication rate under the WC model B) We computed the expected replication rate under the WC+C model

### Application to 100 human GWAS datasets

We then applied our framework to 100 human GWAS previously curated to study Winner’s Curse [4]. All studies have summary statistic data publicly available, a focus on human genetics, and a discovery and replication design, where only the significant SNPs in the discovery study are tested in the replication study. We used the z-scores from these discovery and replication studies as input to our method and estimated the variance parameters (Figure S5, Table S1).

After learning the variance parameters for the genetic and confounding effects, we calculated the estimated replication rates under the two models (Methods, Figure 7). We compared these estimated replication rates to the true replication rates to assess which model explained the observed replication better. We defined the true replication rate to be the proportion of variants in the discovery study that are also significant in the replication study with the same direction of effect in both studies. We used a nominal adjusted p-value threshold of *α* = 0.05 for each replication study. Of the 1652 reported GWAS variants, only 726 (44%) replicated. Using the naive model that does not account for confounding, we would expect 973 (56%) of the variants to replicate. However, when we account for both Winner’s Curse and confounding in our framework, we would expect 762 (46%) of the variants to replicate, which is very close to the observed value.

While the naive model that only accounts for Winner’s Curse explained the replication data well in some cases, in others, we observe a substantial bias beyond what we would expect from statistical noise due to Winner’s Curse (Figure S6). We observed a wide range in the estimated values for the variance in the confounding effects, relative to the variance in the genetic effects (Figure S5). To assess the relative contributions of genetics and confounding to replication, we computed the proportion of variance in the discovery z-scores explained by genetics and confounding (Methods, Equation 22 and Equation 21). We observed a wide range of estimated confounding levels across the 100 studies (Figure 6).

**Figure 6:**
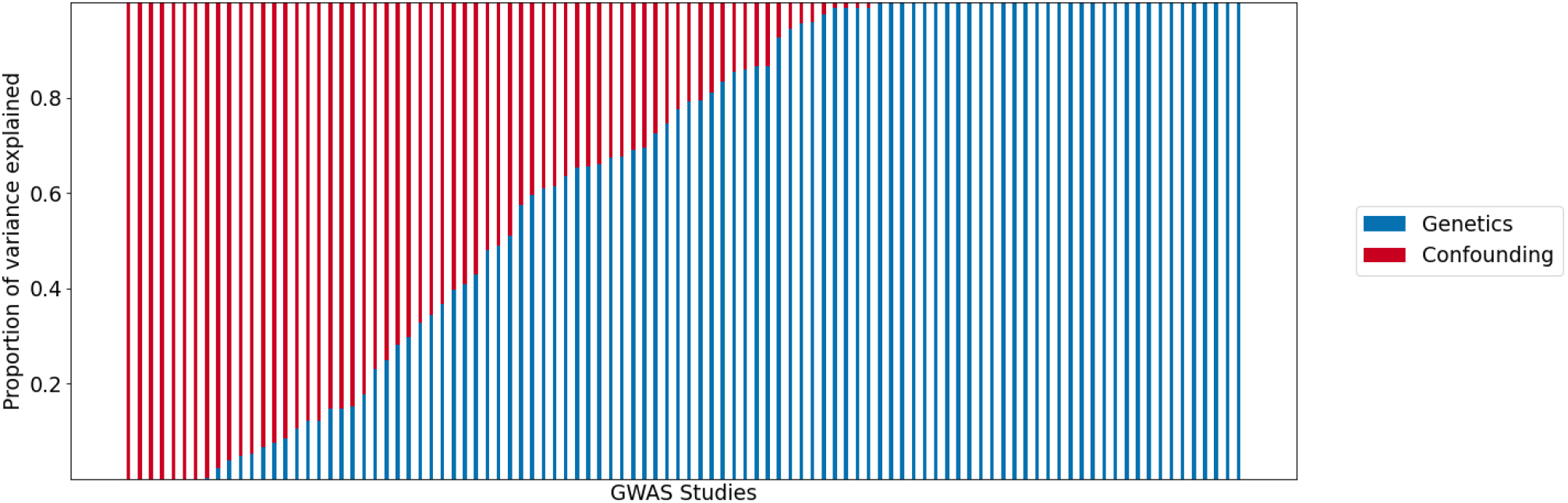
Proportion of variance explained by confounding in 100 human GWAS. Each study is on the x-axis. The proportion of variance explained by genetics (blue) and confounding (red) are shown on the y-axis.

**Figure 7:**
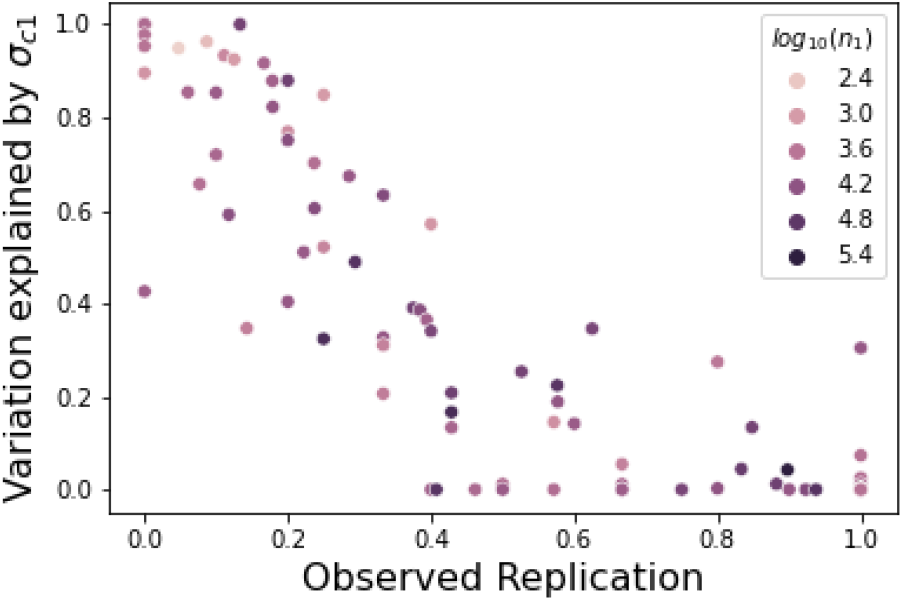
Estimated confounding explains observed replication. The x-axis is the observed replication, and the y-axis is the proportion of variance in the discovery study explained by confounding. Each dot represents a single GWAS study. The Pearson correlation between the estimated variance of confounding and true replication rate is -0.90. The color corresponds to the number of individuals in the discovery study. While the estimated confounding in the discovery study explains the replication rate well, sample size does not explain the replication consistently.

The proportion of variance explained by confounding for the discovery study was strongly correlated (Spearman *ρ* = −.90) with the observed replication, indicating that higher levels of estimated confounding leads to lower replication (Figure 7). However, sample size was not highly correlated with observed replication (Spearman *ρ* = .11) and explained replication inconsistently. While theoretically studies with larger sample sizes tend to have higher power and are more likely to replicate, in practice, some studies with large sample sizes replicate well and others do not.

Similarly, some of the smallest studies had the highest replication rates. Another potential cause of poor replication is noise in the measurement of phenotypes. For example, behavioral phenotypes are often more difficult to measure than physiological traits. While the behavioral phenotypes in this analysis tended to have more confounding than physiological traits, the physiological traits had a wide range in observed confounding levels, indicating that type of phenotype measured also cannot fully explain replication patterns (Figure S7).

Our model can be used to differentiate between cases when lack of replication is due to statistical chance and when it is due to confounding in the data. For example, study 1 (PMID 20935629) is a GWAS on waist hip ratio with 77,167 individuals. The proportion of variance explained by confounding was very low (0%), and a relatively large proportion of variants replicated (94%). This indicates that confounding and statistical noise due to Winner’s Curse did not hinder replication substantially. On the other hand, study 2 (PMID 19079260) is a GWAS on weight and body mass index (BMI), similar phenotypes to waist hip ratio. In this study, only 41% of significant variants replicated. The proportion of variance explained by confounding was still low (0%). Thus, the replication was likely hindered by the smaller sample size (31,392 individuals) and statistical noise contributing to Winner’s Curse. Finally, study 3 (PMID 23669352) is a GWAS on BMI with 29,880 individuals, a similar number of individuals as study 2. However, in this study, only 18% of significant variants replicated. In this study, the estimated level of confounding in the discovery study was very high (82%), which indicates that replication in this study was further hindered by study-specific confounding.

### Comparison to existing corrections of Winner’s Curse

We compared our estimated replication rates under the Winner’s Curse model with those previously reported in Palmer, et al. [4], which corrected for Winner’s Curse using a previously published method, which we refer to as “ZhongPrentice” [23]. At a nominal significance level of 0.05 for the replication study, Palmer et al. estimated that 888 loci would replicate, which is more than the observed replication rate (726 variants). However, it is substantially closer to the observed replication rate than our Winner’s Curse only model, which estimated that 973 variants would replicate.

The primary difference between our estimated replication under the naive Winner’s Curse model and the estimated replication using ZhongPrentice is that our framework model’s Winner’s Curse by accounting for uncertainty in the true effect sizes of the variants. ZhongPrentice treats the true effect size as fixed and attempts to estimate the true effect size by removing the bias due to Winner’s Curse, which is modeled as a function of the true effect and the significance threshold for the discovery study. In this framework, variants with true effect sizes close to the significance threshold of the discovery study have high bias due to Winner’s Curse, regardless of whether the estimated effect size was inflated or not.

In practice, the true effect size is not known, so it is difficult to compare these Winner’s Curse corrections in real data. To compare these two models of Winner’s Curse, we simulated GWAS z-scores for discovery and replication cohorts, where the true effect size was known and the study-specific confounding was set to zero. For each simulation, we fixed *σ*_*g*_ to be one of four values, and we fixed *σ*_*c*1_ = 0 and *σ*_*c*2_ = 0. We simulated z-scores for 1 million independent variants in each simulation. For each combination of parameter values, we repeated this simulation procedure 1000 times to generate a total of 4000 simulations. We then used a Bonferonni threshold of 5e-8 to identify variants that were significant in the discovery study. The number of significant variants in the discovery study for each simulation ranges from 0-1234 variants (Figure S8A). We computed the replication rates as the proportion of variants in the discovery study that met a nominal threshold of 0.05 in the discovery study and had the same direction of effect in the two studies. The observed replication ranged from 0% -84% (Figure S8B).

We computed the MLE estimates of the variance components. For all simulations, the estimation of the parameters was accurate (Figure S9). We computed the difference in the observed and expected replication rates after accounting for Winner’s Curse under our model and ZhongPrentice. As *σ*_*g*_ increases, the difference between the observed and expected replication rates decreases on average for both models (Figure 8). While our method is unbiased for all values of *σ*_*g*_, ZhongPrentice underpredicts the observed replication as *σ*_*g*_ increases(Figure 8).

**Figure 8:**
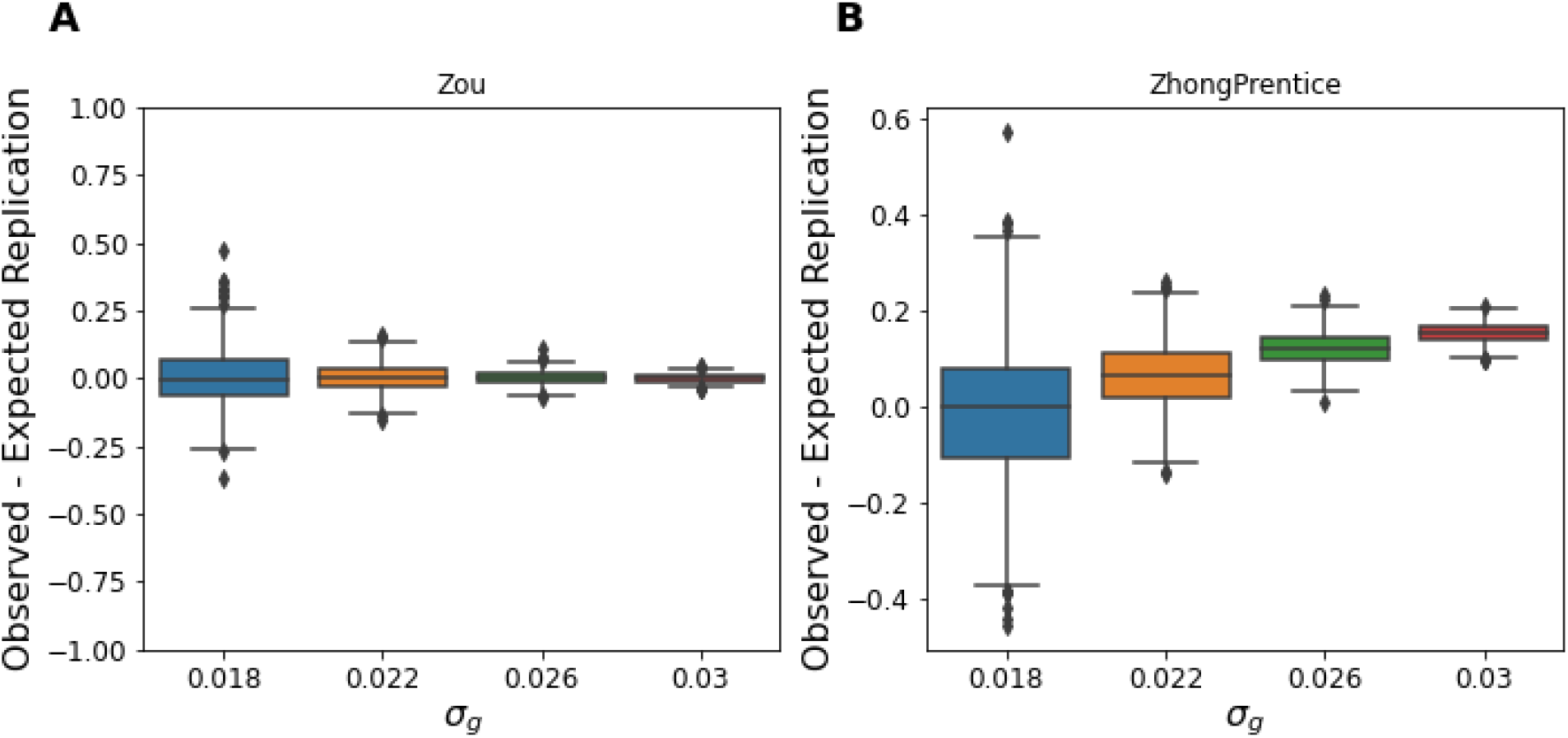
Comparison of methods accounting for Winner’s Curse. We simulated discovery and replication z-scores without study-specific confounding effects for a range of *σ*_*g*_ values. The difference between the observed and expected replication under A) our method B) ZhongPrentice method.

## Discussion

We developed a novel statistical framework to correct for Winner’s Curse and study-specific confounding in GWAS data. This framework utilizes GWAS replications to identify the presence of confounders without relying on assumptions to distinguish between polygenicity and confounding.

We showed through simulations that our model that accounts for Winner’s Curse and Confounding provides accurate estimates of the expected replication rate, even when using incomplete data in a discovery and replication data set. However, the variance in the estimates is higher when using incomplete data to estimate the variance parameters. Thus, if z-scores for variants that are not genome-wide significant are available, it is best to include those variants when estimating the parameters.

When applying our method to 100 human GWAS, we showed that a model that accounts for Winner’s Curse and confounding explains replication rates more accurately than a naive model that only accounts for Winner’s Curse. We observed a range of confounding levels in the 100 GWAS studies analyzed and showed that estimated variance explained by confounding in the discovery study explains the observed replication across studies well, while other factors such as sample size and type of phenotype did not fully explain observed replication.

We demonstrate that our framework can be used to differentiate when studies fail due to statistical noise contributing to Winner’s Curse and when they may fail due to confounding between the studies. One application of this framework would be to identify which studies to include in a meta-analysis or mega-analysis. In GWAS, meta-analysis has discovered many associations that were not identified by each individual study [24, 25]. However, if confounding exists between studies, novel variants found when combining the data could be false positives. It has been proposed to only apply meta-analysis between GWAS that have a high genetic correlation (*rG* > 0.7) [26, 27]. However, it has also been observed that studies can have confounding and poor replication despite high genetic correlation [28]. Our method can be used to determine whether study-specific heterogeneity due to confounders exists before combining data from independent cohorts.

## Methods

### GWAS overview

In GWAS, an association study is performed between each genetic variant and the phenotype. The effect size of each variant (*k*) is determined by estimating the maximum likelihood parameters of Equation 5, where *y*_*j*_ is the phenotype or individual j, *µ* is the phenotypic mean, *x*_*kj*_ is the standardized genotype of variant *k* in individual *j, β*_*k*_ is the effect size of the variant *k, e*_*j*_ is the error, and *N* is the number of individuals.

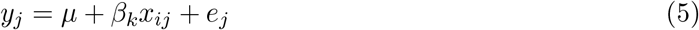

In vector notation, Equation 5 becomes the following.

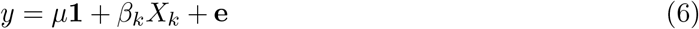

The resulting maximum likelihood estimates are 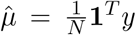 and 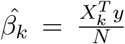. The residuals 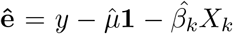 can be used to estimate the standard error 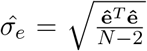. The standard error of the estimator is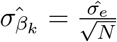. Since the sample sizes for GWAS are large, the association statistic 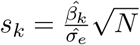 follows an approximately normal distribution (Equation 7).

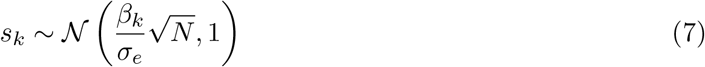

Under the null hypothesis, *S*_*k*_ will follow the standard normal distribution, which can be used to compute the significance of association. In the standard GWAS framework, we assume that the standardized effect size is caused by a true genetic effect 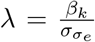. Thus, Equation 7 can be rewritten as the following.

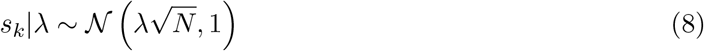

### Correcting GWAS statistics for Winner’s Curse

Let *N*_1_ be the sample size of the discovery study and *N*_2_ be the sample size of the replication study. Given Equation 7, we can write the distributions of association statistics for a discovery study and a replication study as 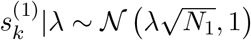 and 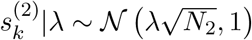, respectively.

We assume that *λ* is the same across multiple studies on the same trait. We define the prior distribution of *λ* as 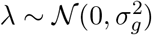, where 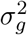 is the variance in the true effect size. Thus, the posterior distributions of 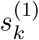 and 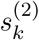 are also normally distributed.

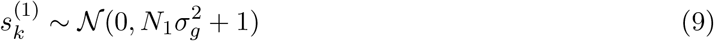

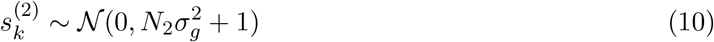

We correct for Winner’s Curse by computing the conditional distribution of the replication statistic 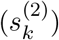 given the discovery statistic 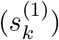. We derive the conditional distribution from the joint distribution as follows.

The covariance between 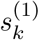 and 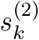 is computed as follows.

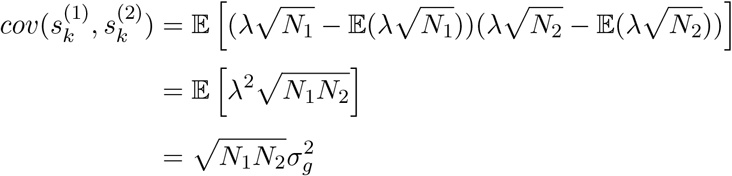

Therefore, the joint distribution of 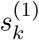 and 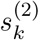 is Equation 11.

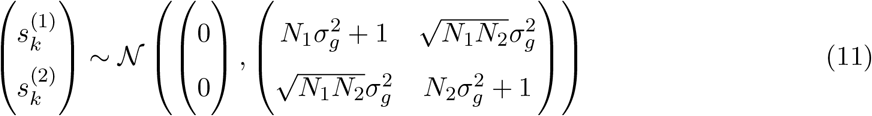

Conditioning on 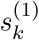, we obtain Equation 12.

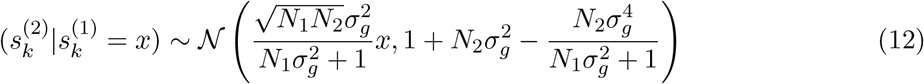

For each value of 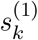, the mean of the conditional distribution gives the expected statistic in a replication study, correcting for Winner’s Curse. This distribution can also be used to create a confidence interval on the replication sample statistics.

### Correcting GWAS statistics for Winner’s Curse and confounding

Suppose in addition to study-specific environmental effects, there are also study-specific confounders. We model these confounders in the discovery study and replication study as 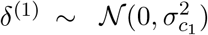 and 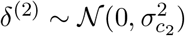 respectively. We decompose the effect size into the sum of a genetic component (*λ*) and a confounding component *δ*^(*i*)^.

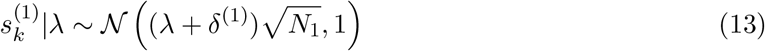

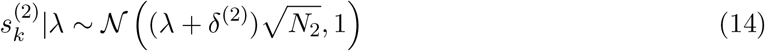

Similar to the case without confounding, the posterior distributions of 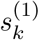 and 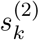 are normally distributed (Equations 15 and 16).

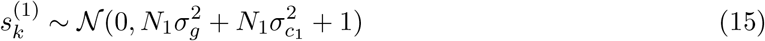

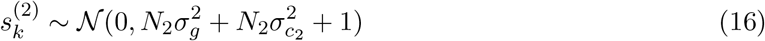

Therefore, the joint distribution is Equation 17

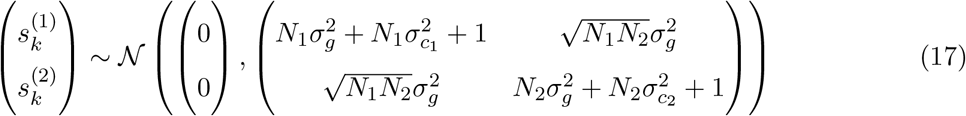

Similar to the Winner’s Curse only model, we can find the expected statistic in a replication study correcting for Winner’s Curse by computing the conditional distribution of the replication statistic 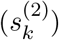 given the discovery statistic 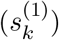 (Equation 18).

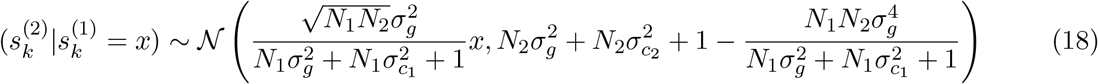

### Estimating the variance components from data

The variance in the true genetic effect 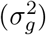 and variance in the confounding effects 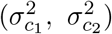 are not known a priori. We estimate these parameters from the data by maximizing the joint likelihood of the discovery and replication z-scores (Equation 17). We compute the maximum likelihood estimators of the variance parameters using the Nelder-Mead method implemented in the scipy.optimize package.

Since typically only a subset of the data that was significant in the discovery is observed, we account for missing data by integrating over all possible values. Let the significance threshold of the discovery study be *t*, and let *z* be the corresponding z-score. We use the joint distribution of the z-scores to compute the probability of a variant not being significant in the discovery study (1 − 2*P* (*s*^(1)^ < *z*)). If the total number of variants that were tested is *N* and the set of significant variants in the discovery study is 𝒜, the negative log-likelihood accounting for missing data is Equation 19.

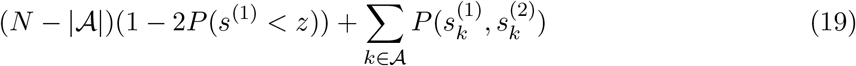

An implementation of our framework is publicly available (https://github.com/jzou1115/wcrep).

### Computing expected replication rates

We computed the expected replication rate under each model using two conditional distributions 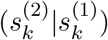 (Equations 18 and 12).

Let 𝒜 be the set of variants found to be significant in the discovery study. We used a nominal threshold of 0.05 for the replication study. Let *z* be the z-score threshold corresponding to *t*. For a genetic variant *k* with association statistic 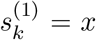 in a discovery study, the probability of replication is 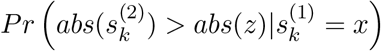. We defined the expected replication rate for a study (*r*) as the average probability of replication for variants significant in the discovery study (Equation 20).

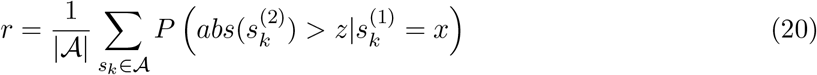

We used the marginal distribution of the discovery summary statistics (Equation 13) to compute the relative proportion of variance explained by genetics and confounding.

We computed the variance explained by genetics *p*_*g*_ as

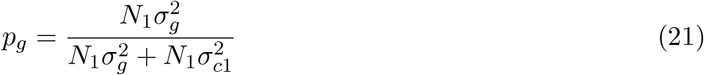

We computed the variance explained by confounding in the discovery study *p*_*c*1_ as

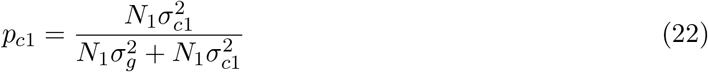

### Data generating model

For all simulations, we fixed the sample size of the discovery study (*N*_1_ = 2000) and the sample size of the replication study (*N*_1_ = 1000).

For each simulation, We fixed the variance parameters to be one of four values *σ*_*g*_, *σ*_*c*1_, *σ*_*c*2_ ∈ [.018, .022, .026, .03]. These values were selected to obtain a realistic range of numbers of significant variants in the discovery study (< 1% of the variants). We simulated summary statistics for 1 million SNPs using the following procedure. For each SNP *k*, we drew true genetic 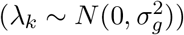 and confounding effects 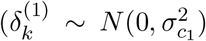 for the discovery study and 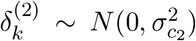 for the replication study). Then, we simulated the z-scores for SNP *k* as the sum of the genetic effect and the study-specific confounding effect, scaled for the sample size of the study.

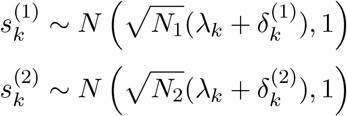

We simulated data for every possible combination of parameter values (4^3^ = 64 combinations) and repeated this procedure 1000 times for a total of 64,000 simulations. For all simulations, we used a Bonferonni corrected threshold of 5*e* − 8 to identify SNPs significant in the discovery study. The observed replication rate was computed as the fraction of variants significant in the discovery study that met a nominal threshold of .05 in the replication study.

We used these simulations to assess the accuracy of our MLE parameter estimates and the expected replication rate under the models under two scenarios: 1) using complete data and 2) using incomplete data. When using complete data, we used the z-scores for all 1 million variants simulated to estimate the variance components. When using incomplete data, we used z-scores for only the variants that were significant in the discovery study.

In order to compare our WC model to previous methods, we generated a second set of simulations. These simulations were identical to the previous set of simulations, except that we fixed the variance in the study-specific confounders to be zero. Thus, we simulated the z-scores for each SNP *k* as

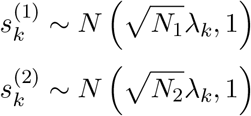

## Acknowledgments

J.Z. and E.E. are supported by National Science Foundation grants 1910885, and 2106908 and NIH grant R56-HG010812. J.Z is supported by a National Science Foundation Graduate Research Fellowship under Grant DGE-1650604.

## Supplementary Materials

Table S1: **Application to 100 human GWAS** We applied our method 100 human GWAS data sets previously published in the articles referenced by PMID. The sample size of the discovery study is “n1” and the sample size of the replication study is “n2”. The significance threshold used in the discovery study is “t”, and the replication threshold is 0.05. The estimated values of the parameters are “sigma g”, “sigma c1”, and “sigma c2”. The total number of variants significant in the discovery study is “num sig”. The number of variants that replicated is “num rep”. The number of variants that are expected to replicate under the WC and WC+C models are “rep wc” and “rep wcc”, respectively. The proportion of variance explained by genetics and confounding are “var exp g” and “var exp c1”, respectively.

**Figure S1:**
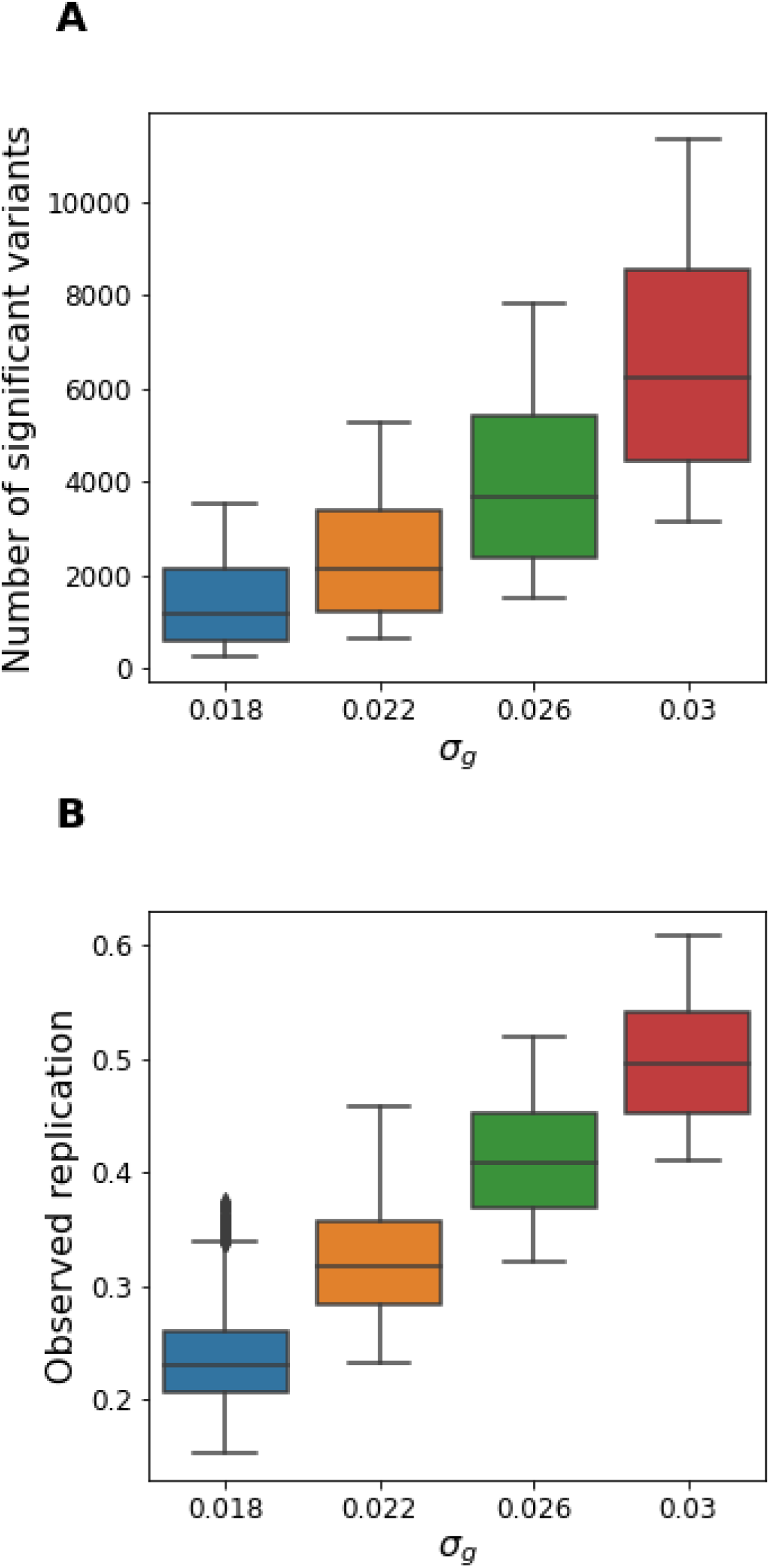
Summary of simulated data. A) Number of significant variants in the discovery study for Winner’s Curse and confounding simulations. We fixed the values of the variance parameters and simulated z-scores for discovery and replication cohorts. The x-axis corresponds to the value of *σ*_*g*_ used to generate the simulations, and the y-axis corresponds to the number of significant variants in the discovery study using a Bonferroni threshold of 5e-8. B) Replication in Winner’s Curse and confounding simulations. We computed the replication rates as the proportion of variants in the discovery study that met a nominal threshold of 0.05 in the discovery study and had the same direction of effect in the two studies.

**Figure S2:**
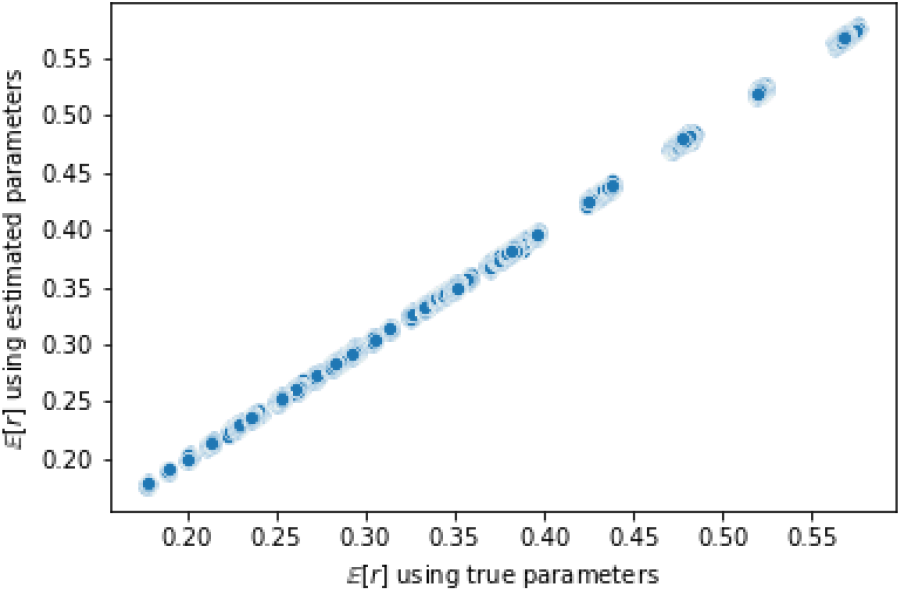
Expected replication rate is robust to variance in MLE parameter estimates. We computed the expected replication rate using the MLE parameter estimates (y-axis) and compared this to the expected replication rate using the true parameters (x-axis). The expected replication using the two sets of parameters are nearly identical, indicating that the parameters are estimated accurately.

**Figure S3:**
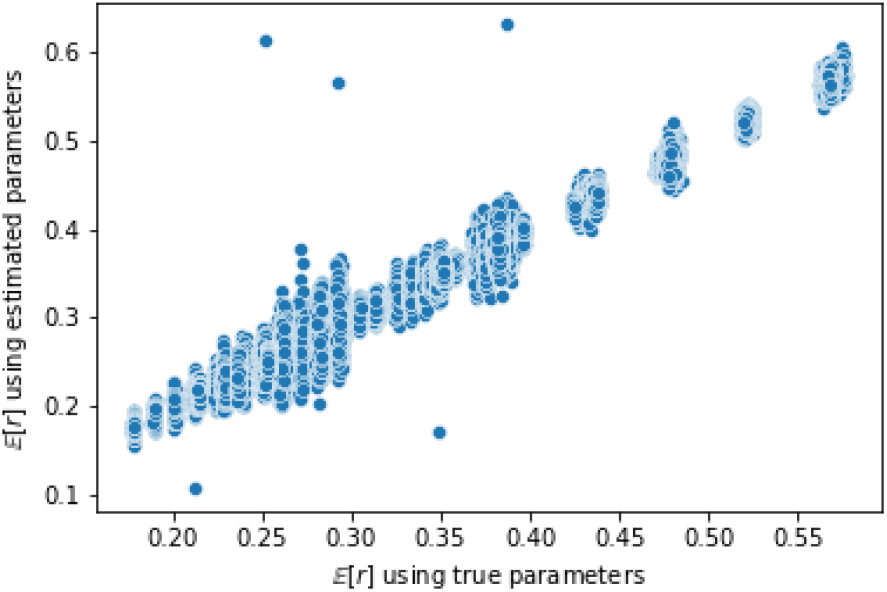
Expected replication rate is robust to variance in MLE parameter estimates with missing data. We computed the expected replication rate using the MLE parameter estimates (y-axis) and compared this to the expected replication rate using the true parameters (x-axis). Despite the increased variance in the parameter estimates when using incomplete data, the expected replication with MLE parameters is similar to expected replication using true parameters.

**Figure S4:**
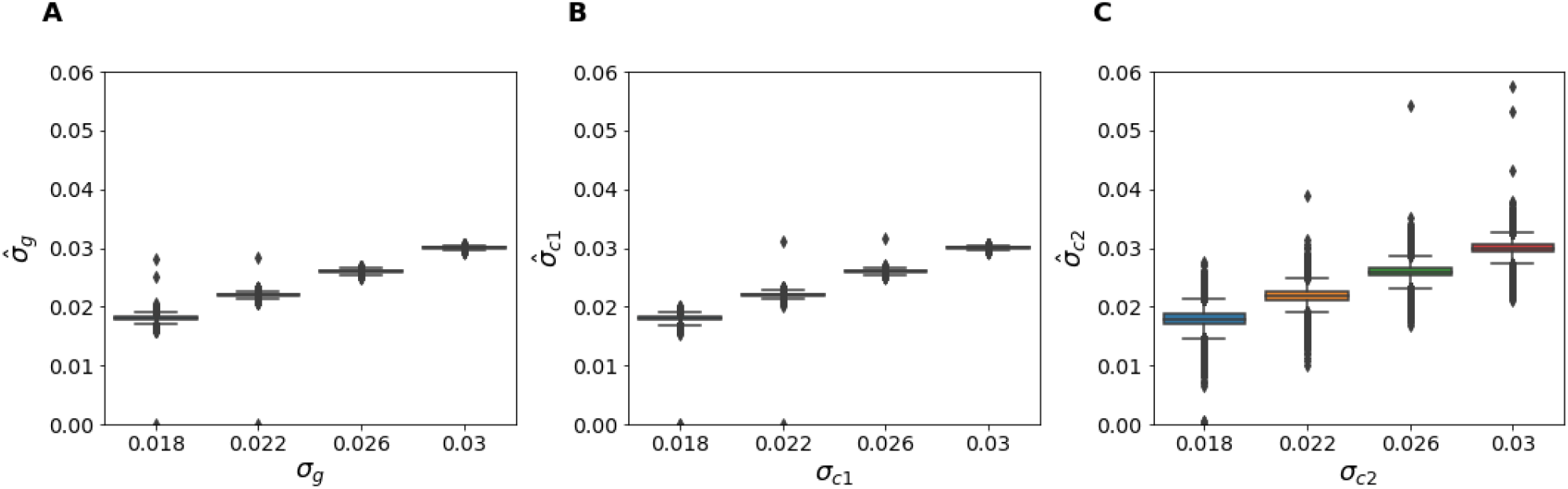
Variance components in Winner’s Curse and Confounding simulations with incomplete data. True values of variance components (x-axis) vs estimated values (y-axis) for (A) 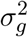 B) 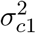 C) 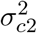

**Figure S5:**
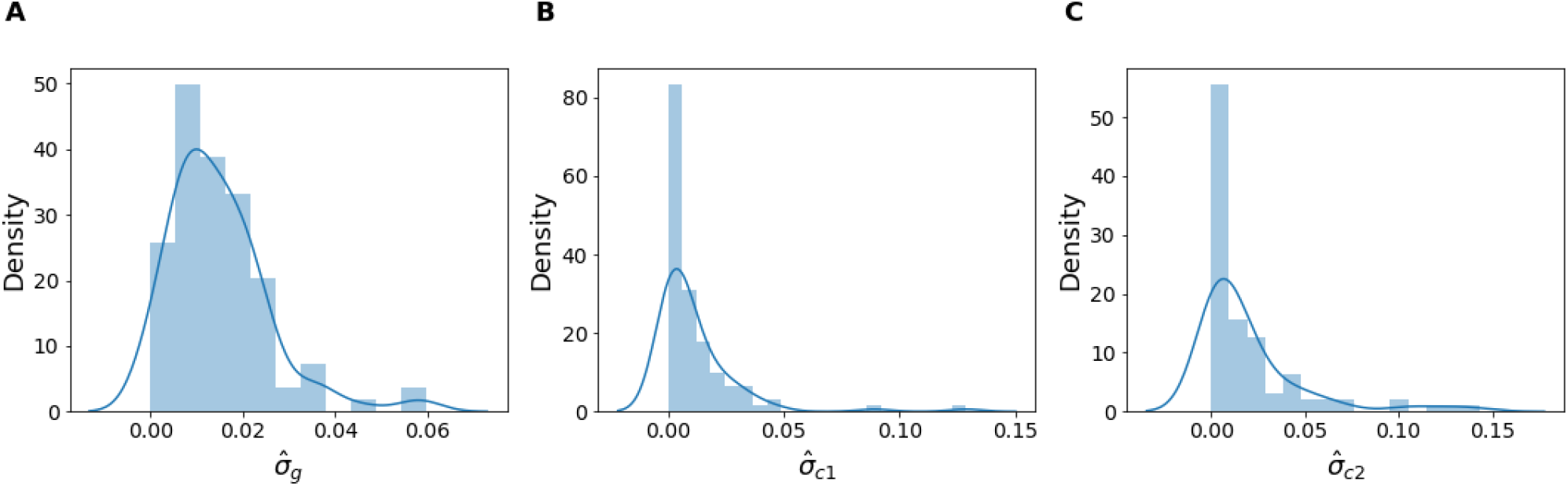
Distribution of variance components in 100 human GWAS. Distribution of MLE estimates of A) *σ*_*g*_ B) *σ*_*c*1_ c)*σ*_*c*2_

**Figure S6:**
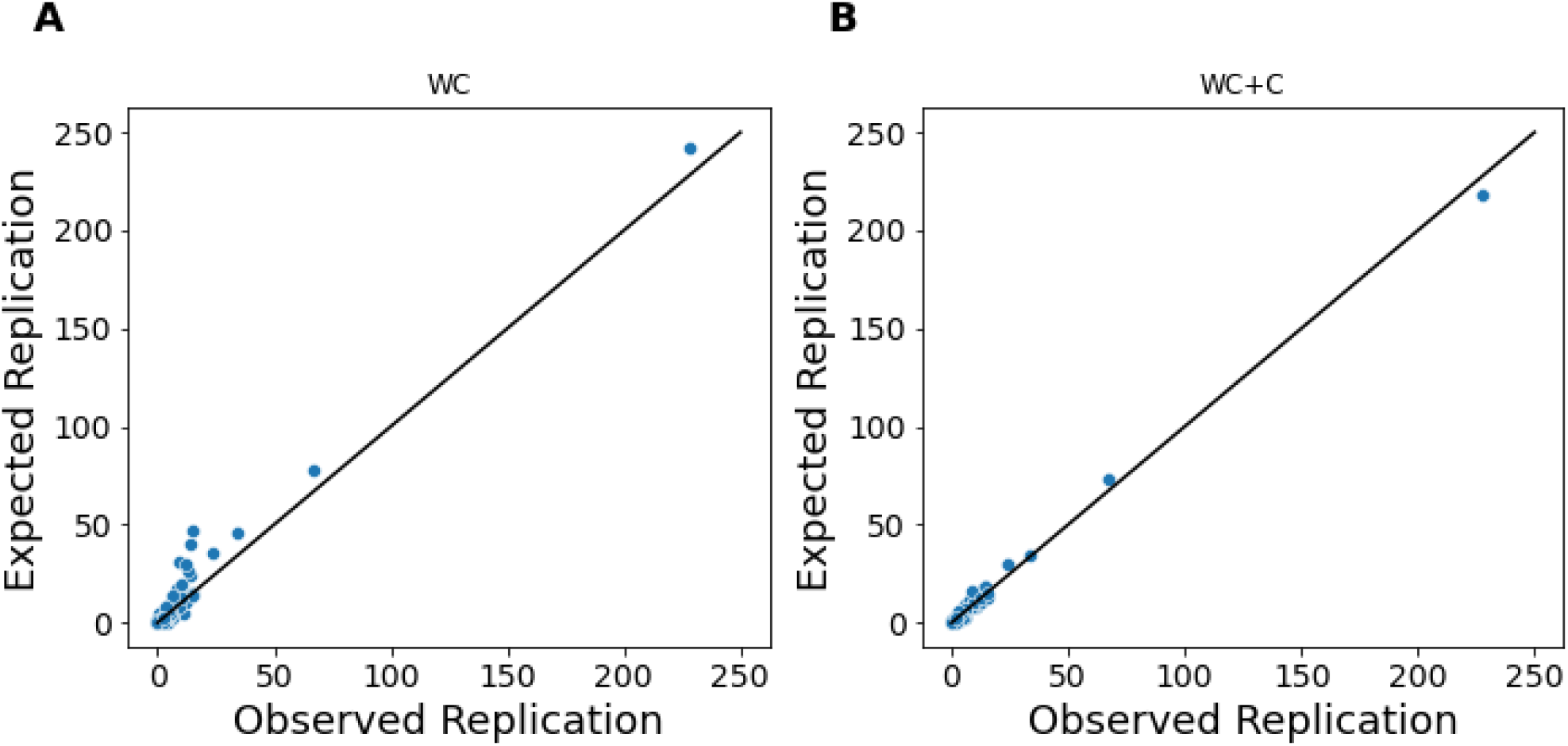
Expected replication in 100 human GWAS. The x-axis is the true number of variants that replicate. The y-axis is the estimated number of variants that replicate under a model. Each dot represents one GWAS study. A) Expected replication under WC model B) Expected Replication under WC+C model

**Figure S7:**
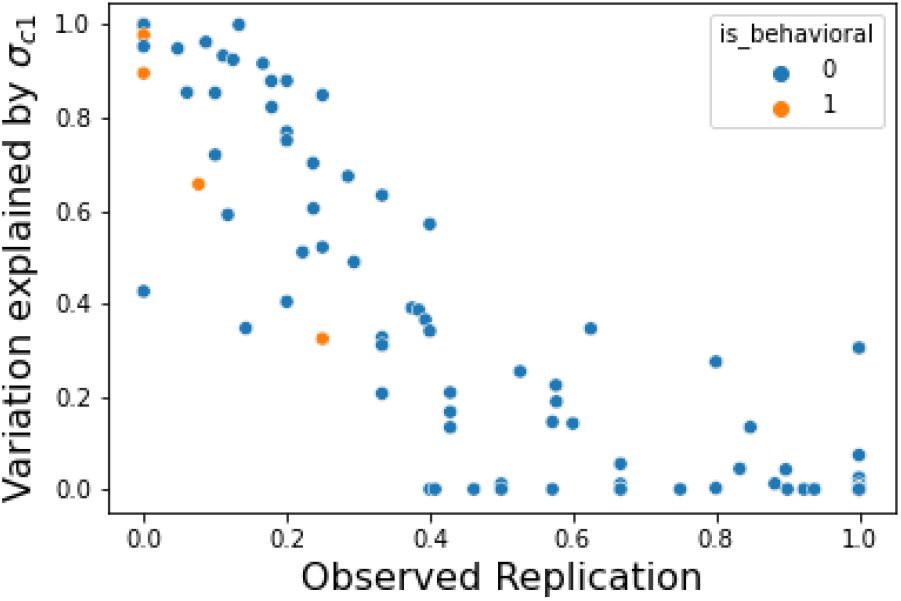
Behavioral phenotypes have higher levels of confounding. The x-axis is the MLE of 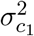, and the y-axis is the true replication rate. Each dot represents a single GWAS study. The color corresponds to whether the phenotype is behavioral or not. Behavioral phenotypes tend to have higher estimated levels of confounding.

**Figure S8:**
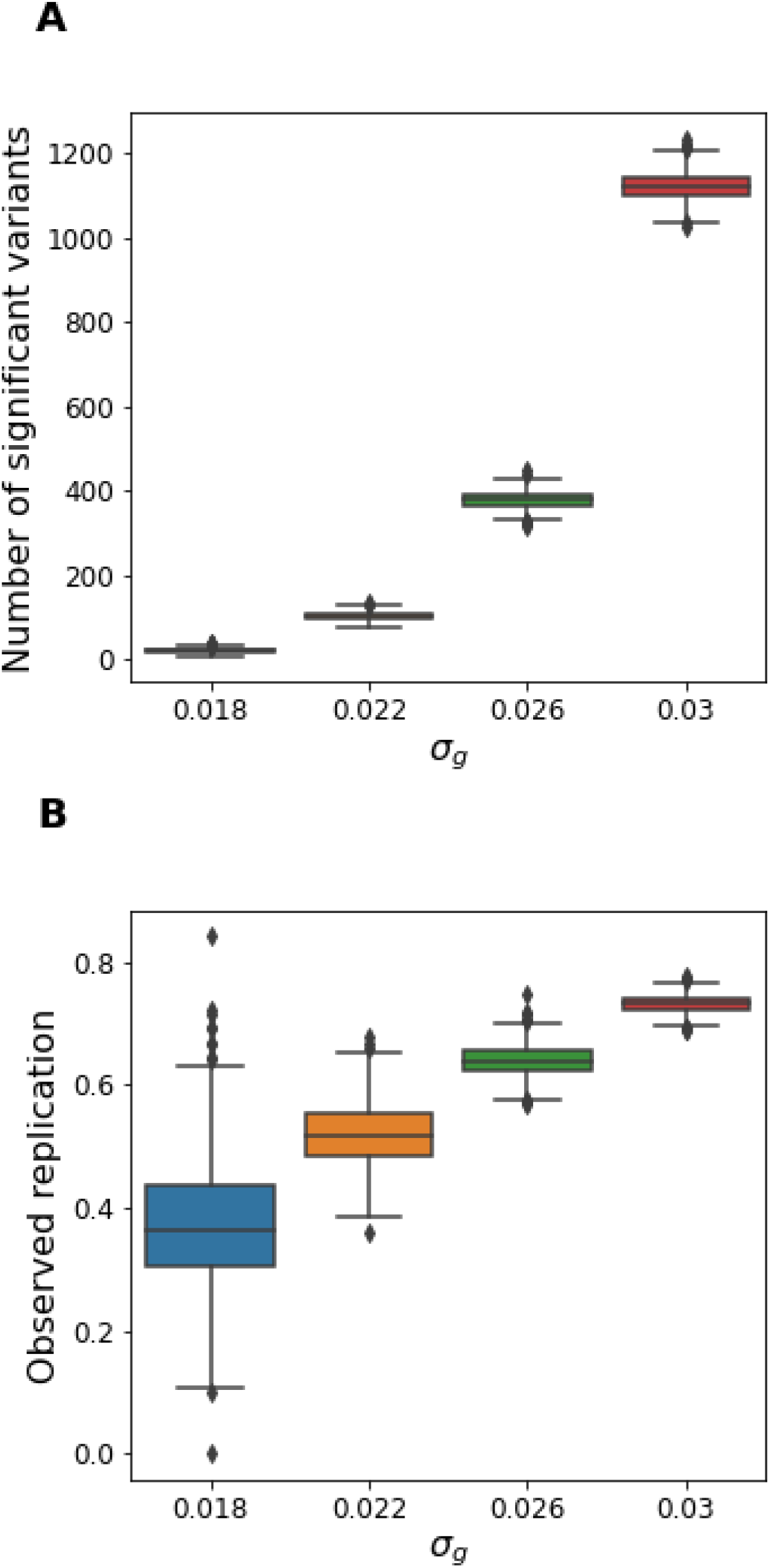
Summary of simulated data for Winner’s Curse comparisons. A) Number of significant variants in the discovery study for Winner’s Curse simulations. We fixed the values of the variance parameters and simulated z-scores for discovery and replication cohorts. The x-axis corresponds to the value of *σ*_*g*_ used to generate the simulations, and the y-axis corresponds to the number of significant variants in the discovery study using a Bonferroni threshold of 5e-8. B) Replication in Winner’s Curse and confounding simulations. We computed the replication rates as the proportion of variants in the discovery study that met a nominal threshold of 0.05 in the discovery study and had the same direction of effect in the two studies.

**Figure S9:**
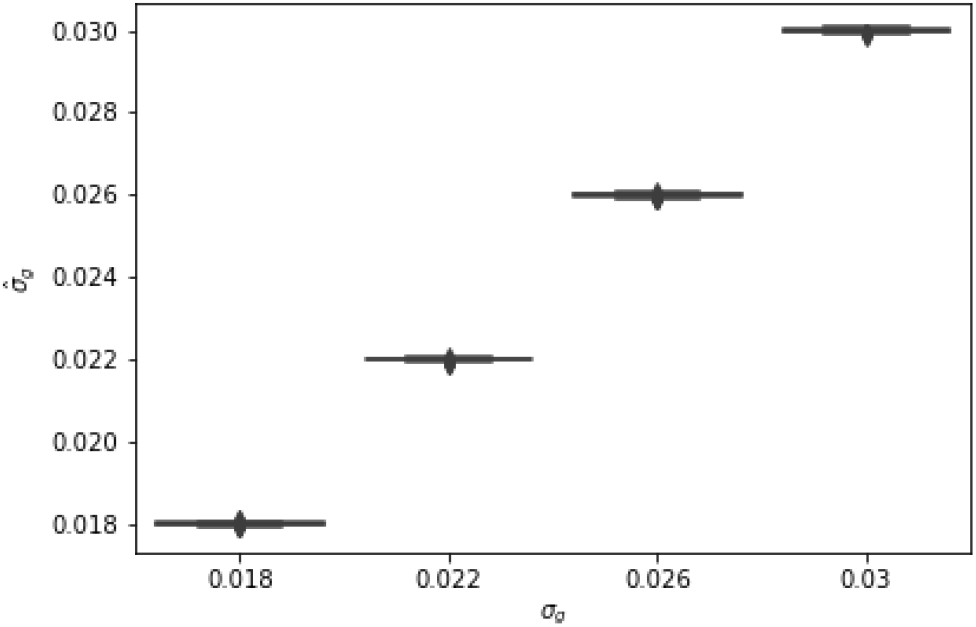
Variance Components in Winner’s Curse Simulations. True values of variance components (x-axis) vs estimated values (y-axis) for 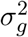

